# Intervertebral disc impairments in a mouse model of Alzheimer’s Disease

**DOI:** 10.1101/2025.05.24.655931

**Authors:** CE Gonzalez, CA Schurman, KA Wilson, B Schilling, LM Ellerby, SY Tang

**Affiliations:** Department of Biomedical Engineering, Washington University in St Louis, St. Louis, MO; Department of Orthopaedic Surgery, Washington University School of Medicine, St Louis, MO; Buck Institute for Research on Aging, Novato, CA, USA

**Keywords:** Alzheimer’s disease, intervertebral disc degeneration, 5xFAD, amyloid dysregulation, chronic low back pain

## Abstract

Chronic low back pain, frequently associated with intervertebral disc (IVD) degeneration, is highly prevalent in individuals with Alzheimer’s disease (AD), and the pain intensity is highly correlated with the degree of cognitive impairment. While the incidences of both afflictions increase dramatically in the elderly population, it is unknown whether AD exacerbates the health of the IVD. Utilizing one-year-old male and female 5xFAD mice that constitutively express human APP and PSEN1 transgenes with five AD-linked mutations, we measured the lumbar IVD’s extracellular matrix composition, the three-dimensional structure, histopathological degeneration, and mechanical behavior. The collagen, glycosaminoglycans, and advanced glycation end-products content of the IVD were not appreciably different between the 5xFAD animals and their wild-type littermates. Likewise, the 5xFAD IVDs were not histopathologically degenerated. However, the IVD volume, measured by contrast-enhanced microCT, was larger in the 5xFAD animals. Furthermore, dynamic microcompression revealed that 5xFAD IVDs exhibited higher loss tangent, indicating altered tissue damping and fluid-flow dynamics within the disc. These results suggest that although the IVDs of mice with AD are not more degenerated, they may be more susceptible to damage accumulation due to the elevated absorption of energy. Elderly individuals with AD may thus be more prone to IVD injuries that lead to eventual degeneration and spinal pain. Future work will focus on defining the molecular mechanisms and the consequences of these mechanical and structural changes in the IVD and their consequences to low back pain in individuals with AD.

## Introduction

Alzheimer’s disease (AD) is a progressive neurodegenerative disorder characterized by cognitive decline, memory impairment, and behavioral changes, and it significantly deteriorates the quality of life in patients. Concurrently, chronic low back pain (cLBP) is also increasingly prevalent among older adults, with recent data indicating that elderly individuals are experiencing cLBP at a more than two-fold increase over the past decade (Weiner, 2015). The etiology of cLBP involves multifactorial mechanisms including degenerative changes in spinal tissues, inflammation, and biomechanical stress. Emerging evidence suggests a critical association between cLBP and AD, proposing shared pathological mechanisms such as neuroinflammation and locus coeruleus noradrenergic dysfunction, which may exacerbate cognitive decline (Cao et al., 2019). This intersection underscores the urgent need for an improved understanding of these mechanisms.

The management of pain in patients with AD presents unique challenges due to impaired cognitive function, leading to difficulties in pain reporting and frequent misinterpretation of pain-related behaviors by clinicians (Malotte and McPherson, 2016; Rababa, 2018). These issues often result in the undertreatment of pain, exacerbating suffering, and complicating disease progression (Belin and Gatt, 2006; Romm, 2023). Given the complexity and variability of pain presentation in this population, it is crucial to identify intrinsic risk factors associated with spinal pain and to better understand the underlying mechanisms of comorbid pain conditions. Specifically, it is imperative to determine whether individuals with AD are particularly susceptible to factors that contribute to cLBP.

Despite recognition of intervertebral disc (IVD) degeneration as a major contributor to cLBP through mechanisms like neurogenic inflammation and nociceptor sensitization, gaps remain in understanding the role of amyloid-driven neurodegeneration in musculoskeletal pain sensitization. IVD degeneration triggers nociceptor activation and inflammatory cytokine release, enhancing pain perception through peripheral and central sensitization (Brisby, 2006; Peng et al., 2025; Chiu et al., 2025; Jiang et al., 2024).

To explore the interactions between amyloid pathology and IVD health, preclinical animal models, notably the 5xFAD mouse strain, provide an invaluable tool for longitudinal studies. The five familial Alzheimer’s disease mutations in the 5xFAD model—three in APP (KM670/671NL, I716V, V717I) and two in PSEN1 (M146L, L286V)—are known to drive high levels of amyloid-beta deposition that is not limited to the central nervous system (Koskela et al., 2025). Already, it has been demonstrated that altered cortical bone architecture present in 5xFAD males animals, suggesting systemic effects possibly mediated by peripheral Aβ or associated inflammatory signaling (Suryadevara et al., 2020). Additionally, PSEN1 mutations can modulate Notch and γ-secretase pathways, which influence ECM remodeling and cell fate in many tissues, including the spine (Hurley et al., 2023). These systemic alterations may disrupt intervertebral disc homeostasis through subtle changes in matrix turnover or mechanical behavior. These models recapitulate the feature of amyloid plaque accumulation in human AD, allowing researchers to investigate systemic musculoskeletal impacts, including early biomechanical and compositional alterations in the spine (Akhtar et al., 2022). Given the limitations inherent in human clinical studies—such as delayed AD diagnosis and cognitive-related underreporting of pain—animal models offer controlled environments to elucidate subtle yet clinically relevant changes. Therefore, we utilized the 5xFAD mouse model to investigate structural and functional changes in lumbar intervertebral discs under conditions of disrupted amyloid pathology.

## Materials & Methods

### Animal model

This study utilized male and female 5xFAD heterozygous mice and wild-type (WT) littermate controls, aged 12 months, with n=8 per group (5 males and 3 females) in a C57/BL6 background (Jackson Laboratories, MMRRC_034848-JAX). After confirming the appropriate genotypes, the animals were allowed to age to 12 months and then euthanized in compliance with animal welfare regulations. The age of 12 months was selected to evaluate the systemic consequences of chronic amyloid burden, corresponding to an advanced stage of Alzheimer’s pathology in the 5xFAD model. By this age, mice exhibit robust and widespread amyloid plaque deposition across cortical and hippocampal regions, accompanied by synaptic loss, gliosis, and behavioral deficits (Oakley et al., 2006; Sasaguri et al., 2017). All protocols and procedures described herein were approved by the Buck Institute’s AAALAC-accredited Institutional Animal and Use Committee (Unit Number 001070).

### Structural and histological assessments

Functional spine units (FSUs) were excised from the lumbar spine (L1/L2 to L5/L6) and divided the following assays. For histological assessment, FSUs were fixed in 10% neutral-buffered formalin overnight. After fixation, samples were decalcified for 3 days in ImmunoCal formic acid solution, then washed in phosphate-buffered saline and dehydrated sequentially with 30%, 50%, and 70% ethanol in water. Samples were then embedded in paraffin blocks and sectioned via microtome at a thickness of 10 µm. Sections were stained with Safranin-O/Fast Green, and slides were imaged using the Hamamatsu NanoZoomer Microscope with a 20x objective. Section images were then blindly graded for histopathological degeneration according to a standardized scoring rubric covering different aspects of IVD degeneration (Melgoza et al., 2021). Finally, excised IVDs were used to measure the biochemical content of various matrix proteins. First, discs were digested overnight in a papain buffer and then the lysates were used for a 1,9-dimethylmethylene blue assay for sulfated glycosaminoglycan content with a chondroitin sulfate standard (Liu et al., 2017). The disc was hydrolyzed at 120°C in 12 N HCl; hydrolysates were allowed to evaporate, reconstituted with 0.1x phosphate-buffered saline, and then measured against a quinine standard for advanced glycation end product content (Liu et al., 2017). Finally, a hydroxyproline assay was used to quantify collagen content as previously described (Liu et al., 2017).

### Imaging and morphological analyses

Functional spine units were incubated in 175 mg/mL Ioversol solution (OptiRay 350; Guerbet, St. Louis) diluted in PBS; after 4 hours in contrast agent, samples were scanned using micro-computed tomography (µCT) (VivaCT40, Scanco Medical AG) at 10-µm voxel size, using 45 kVp, 177 µA, and 300 ms integration. Whole disc volume and mean intensity were calculated from contours drawn in a custom MATLAB program (https://github.com/WashUMusculoskeletalCore/Washington-University-Musculoskeletal-Image-Analyses). Disc height index (DHI) was measured by averaging the height-to-width ratio of the IVD over five slices in the mid-sagittal plane, using the linear measure tool in this program (Lin et al., 2016).

### Mechanical testing

Functional spine units were tested using cyclic compression on the BioDent microindenter (Active Life Scientific) with a 2.39 mm OD probe. Samples were fixed to a metal plate and placed in PBS prior to aligning the sample parallel with the probe under 0.03 N preload. Each unit was then sinusoidally loaded at 1 Hz for 20 cycles with a 35 μm amplitude. Loading slope was calculated from the force-displacement curve, the loss tangent (tan delta) was calculated from the loading-displacement delay, and the energy dissipated was calculated by computational integration of the loading hysteresis. Analyses were performed using MATLAB.

### Statistical approach

Statistical analyses were conducted using Welch’s Unpaired t-test in GraphPad Prism 10.4.2. Statistical significance was defined at a threshold of p<0.05. All data were expressed as means ± standard error of the mean.

## Results

### Animal health

The 5xFAD mice were in general good health with relevant phenotypes in the open field and learning and memory consistent with prior descriptions (Makhijani et al., 2024).

### Matrix composition of the IVD is not significantly disrupted in 5xFAD mice with no notable histopathologic degeneration

Histopathological analyses revealed no significant difference in degenerative score between 5xFAD and WT IVDs, confirming preserved gross structural integrity across both groups (Fig. 1A, 1B). Biochemical quantifications for collagen, glycosaminoglycan (GAG), and advanced glycation end-products (AGE) content showed no statistically significant differences, demonstrating that these majority components of the extracellular matrix are not altered by systemic alterations to amyloid metabolism (Fig. 1C–E).

**Figure 1.**
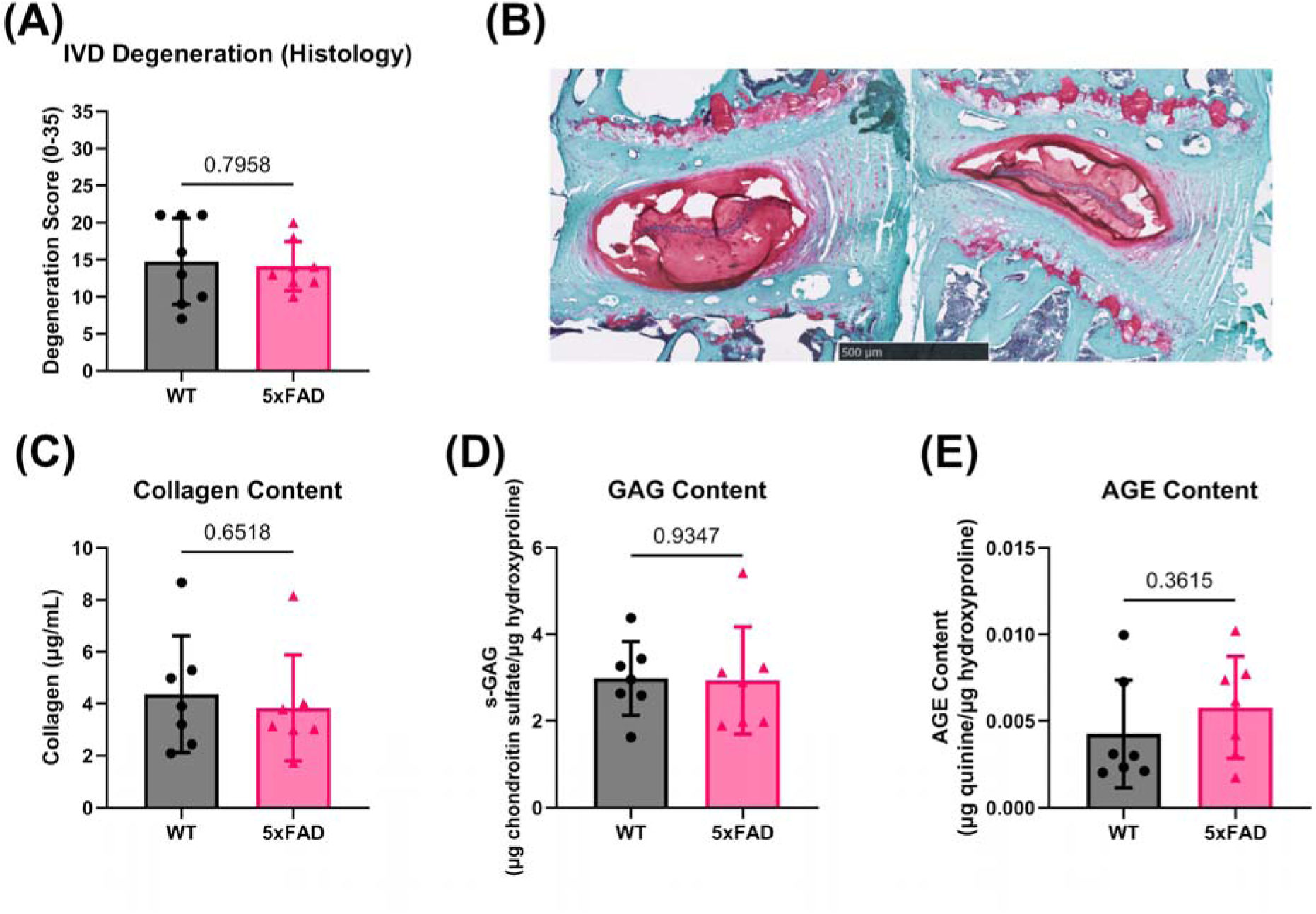
Histopathological and biochemical assessments of IVDs reveal that there no differences in gross tissue degeneration and major components of the extracellular matrix. **(A)** No statistical difference in histopathological grades of IVD degeneration were detected between WT mice and 5xFAD mice. Grading covers the different components of the IVD - nucleus pulposus, annulus fibrosus, endplate, boundary (NP, AF, EP, Boundary) **(B)** Representative Safranin-O/Fast Green staining images illustrating preserved matrix integrity in both 5xFAD and WT groups. **(C–E)** Quantitative analyses of collagen, glycosaminoglycan (GAG), and advanced glycation end-products(AGEs) demonstrating no statistically significant differences between groups.

### Contrast-enhanced µCT shows 5xFAD IVDs are larger than WT

Imaging analyses indicated no significant changes in disc height index between groups, indicating that disc collapse or overt structural degeneration had not occurred by 12 months of age (Fig. 2A). However, 5xFAD IVDs exhibited a significantly larger total volume compared to WT controls, potentially subtle changes in extracellular matrix distribution (Fig. 2B). Mean CT intensity was lower, although statistically non-significant, in the 5xFAD group, possibly indicative of minor variations in hydration or ECM constituents (Fig. 2C). 3D reconstructions from CT scans confirm the observed quantitative enlargement of the 5xFAD IVDs (Fig. 2D).

**Figure 2.**
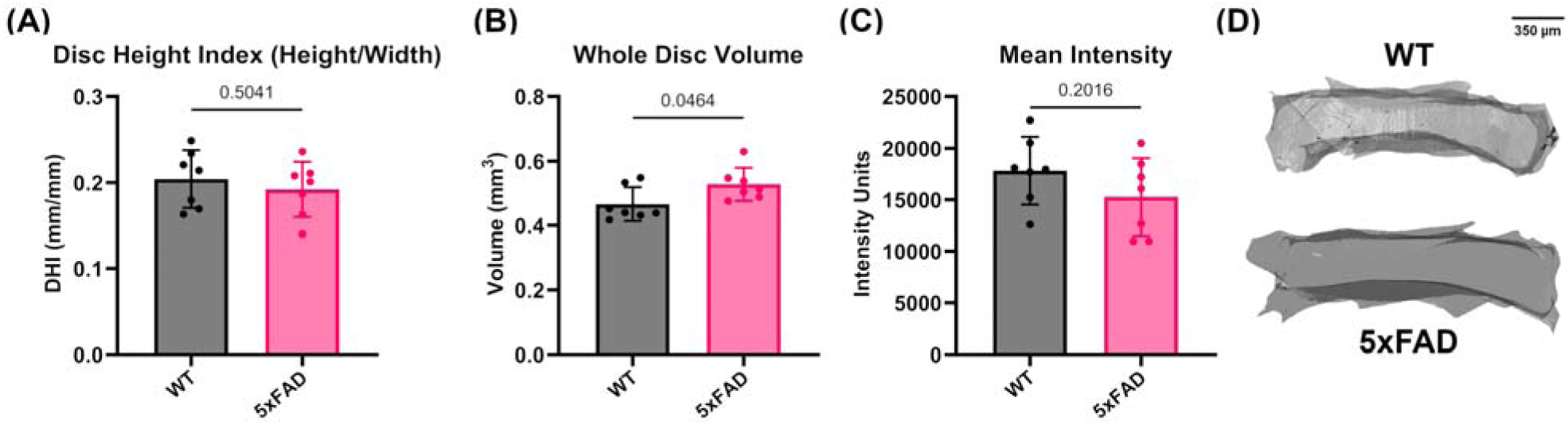
Imaging and morphological analyses of IVDs via contrast-enhanced µCT. **(A)** Disc height index showed no significant differences between groups. **(B)** Quantitative comparison of total IVD volume, highlighting statistically significant enlargement in 5xFAD mice. **(C)** Mean CT intensity, indicating slight but insignificant reduction in 5xFAD discs. **(D)** Representative 3D reconstructions of IVDs confirming quantitative findings.

### Mechanical properties reveal 5xFAD IVDs have higher loss tangent

Mechanical testing revealed a significantly elevated loss tangent in 5xFAD discs (Fig. 3A), suggesting increased viscous behavior compared to WT discs, which might predispose them to mechanical damage under repetitive loading conditions. Energy dissipation showed a trend towards elevation in 5xFAD discs, although this did not reach statistical significance (Fig. 3B). Stiffness measurements did not differ significantly between groups (Fig. 3C), despite minor discrepancies, reinforcing the concept of early subtle mechanical changes rather than extensive structural degradation.

**Figure 3.**
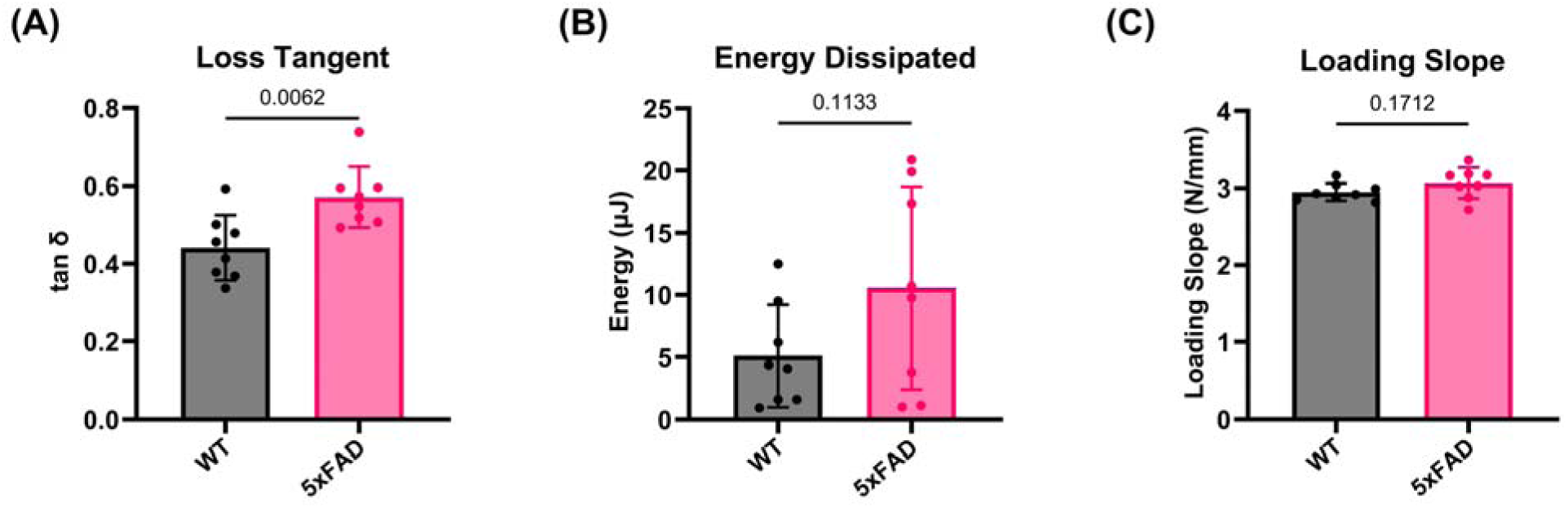
Mechanical testing of IVDs under dynamic compression. **(A)** Loss tangent, significantly elevated in 5xFAD IVDs, indicative of higher viscosity compared to WT controls. **(B)** Energy dissipation, demonstrating a trend toward increased energy loss in 5xFAD discs, although not achieving significance. **(C)** Loading slope indicates stiffness no differences between groups.

## Discussion

The structural and mechanical findings in the present study, characterized by increased IVD volume and elevated viscous mechanical behavior indicate early biomechanical dysfunction preceding overt degeneration, independently of preserved histopathology and composition of the matrix components. These are likely driven by alterations in tissue fluid dynamics and microstructural matrix organization of the IVDs of 5xFAD mice. A shifts towards increased viscous behavior can impair the IVD’s ability to recover from mechanical loading, promoting microdamage accumulation and rendering the IVD more vulnerable to progressive degeneration (Easson et al., 2023; Paul et al., 2018; Sciortino et al., 2024). Such biomechanical impairments potentially stem from modifications at the molecular or ultrastructural level, including altered proteoglycan interactions and disrupted collagen network tension, thereby establishing a mechanistic connection between systemic amyloid pathology and musculoskeletal degeneration.

Functional impairments within IVDs may occur independently of measurable changes in its majority constituents like collagen and glycosaminoglycans. The IVD’s mechanical integrity is highly dependent upon not only matrix composition but also the precise spatial organization and physical interactions of its extracellular matrix components. Proteoglycan-induced osmotic pressures maintain hydration, while collagen networks provide tensile stability (Feng et al., 2006). Subtle disruptions in these microstructural features—such as altered collagen fibril alignment, reduced cross-linking, or changes in proteoglycan-collagen interactions—can adversely affect fluid dynamics and mechanical load distribution without necessarily altering total biochemical content (Colombini et al., 2008). These disruptions can compromise the tissue’s mechanical resilience, shifting the disc’s biomechanical behavior toward increased viscosity, which results in greater energy dissipation and diminished elastic recovery under mechanical stress.

The functional consequences of elevated viscous behavior include reduced effectiveness in load distribution and diminished elastic recovery after deformation. Healthy IVDs temporarily store and gradually release mechanical energy, facilitating efficient load transfer between vertebral bodies. Increased viscosity disrupts this balance, causing prolonged deformation and delayed recovery, leading to localized stress accumulation within disc components (Dougill, 2016). Over time, these stress concentrations can cause microdamage, particularly to collagen fibers and proteoglycan structures responsible for tissue hydration and integrity (Azarnoosh et al., 2014). Consequently, the mechanical resilience of the disc deteriorates further, accelerating degenerative processes and potentially contributing to clinical outcomes such as chronic low back pain.

An increased IVD volume without concomitant biochemical changes suggests altered water retention dynamics driven by subtle organizational shifts in proteoglycan networks. The nucleus pulposus osmotic pressure, largely influenced by proteoglycans, maintains hydration and volume (Adams, 2015). Minor disorganization within the proteoglycan-hyaluronic acid network can increase water content and disc volume, even if glycosaminoglycan and collagen quantities remain stable (Jay Lipson and Muir, 1981). Moreover, they can be caused by changes in minor components, such as decorin, which are not measured here (Sao and Risbud, 2024). This highlights that biomechanical abnormalities may initially manifest from architectural changes rather than outright macromolecular degradation, providing early indicators of potential degenerative progression, especially under systemic pathological conditions like AD.

Limitations of this study include its cross-sectional design, which restricts understanding of temporal changes in intervertebral discs during AD. Additionally, the lack of direct spinal amyloid quantification prevents definitive causal connections between amyloid pathology and IVD dysfunction. Behavioral assessments were not included, limiting insights into functional relevance, such as pain or mobility. The relatively small sample size may also affect the generalizability of findings. Furthermore, the use of a single animal model may not fully capture the complexity of AD pathology observed in humans. Finally, sex-specific differences were not examined, potentially overlooking variations in disease progression or biomechanical responses between males and females. Future studies incorporating longitudinal designs, spinal amyloid measurements, behavioral assays, larger cohorts, multiple animal models, and sex-specific analyses would address these limitations and provide more robust conclusions.

Mechanistically, future research should investigate how amyloid precursor proteins and amyloid-beta may dysregulate extracellular matrix signaling pathways essential for IVD homeostasis. Pathways such as NF-κB and RAGE, potentially influenced by amyloidogenic stress, warrant examination to understand their roles in inflammatory responses and matrix turnover disruptions (Molinos et al., 2015). Additionally, investigating amyloid’s impact on mechanotransduction and cell-matrix adhesion could reveal critical insights into disc degeneration mechanisms and potential therapeutic targets (Mohanty and Dahia, 2019). Aβ, a hallmark peptide in AD, has increasingly been implicated in disrupting these fundamental mechanotransductive processes beyond the central nervous system. In neurons, Aβ disrupts actin and microtubule organization via altered phosphorylation cascades and cytoskeletal destabilization, undermining cellular architecture and impairing signal transduction pathways essential for mechanical responsiveness (Henriques et al., 2015). Similar disruptions have been observed in skeletal muscle and bone. Mukhamedyarov et al. demonstrated that Aβ impairs skeletal muscle excitability by inhibiting Na /K -ATPase and forming membrane-penetrating “amyloid channels,” leading to depolarization and mechanical dysfunction (Mukhamedyarov et al., 2013). In cartilage and bone, mechanosensitive cells like chondrocytes and osteocytes depend on tightly regulated cytoskeletal and receptor-linked pathways to convert strain into anabolic responses; however, Aβ may interfere with these cascades, potentially mimicking effects seen in neurons by attenuating integrin signaling or modulating calcium influx (Goodman et al., 2015; Papachristou et al., 2009). Moreover, recent evidence shows that Aβ directly suppresses TRPV4-mediated mechanosensation in dorsal root ganglion neurons, a channel also implicated in skeletal mechanotransduction (Easson et al., 2023; Fukazawa et al., 2024). These findings suggest that amyloid presence broadly compromises the ability of mechanically active tissues to sense and respond to load. Given the similar mechanobiological reliance of IVD cells, it is plausible that systemic Aβ burden disrupts mechanotransduction within the IVD, leading to altered matrix regulation and mechanical stability.

Clinically, the underreporting of pain in individuals with AD presents a significant barrier to the early detection and effective management of spinal pathologies. Cognitive impairments common in AD interfere with a patient’s ability to recognize, interpret, and communicate pain, which often leads to delays in diagnosis and undertreatment of musculoskeletal conditions such as low back pain (Wang et al., 2022). At the tissue level, emerging evidence suggests that disruptions relevant to AD may influence the biomechanics of the IVD, particularly by increasing viscous behavior—a change that can impair the disc’s ability to distribute mechanical loads efficiently (Liu et al., 2015; Sun et al., 2023). This altered load-bearing capacity, along with cumulative microdamage to the tissue matrix, may make the IVD more vulnerable to degeneration during everyday mechanical stress, predisposing individuals to low back pain (Sciortino et al., 2024). Concurrently, systemic effects of AD may disrupt the mechanotransduction pathways within IVD cells, impairing their ability to sense and respond to mechanical and biochemical cues (Azarnoosh et al., 2014). Such disruptions could affect how the disc maintains homeostasis and interprets internal physiological changes, possibly altering proprioceptive and interoceptive signaling. Finally, changes in central nervous system processing associated with AD can further distort pain perception, reducing the likelihood that spinal discomfort will be noticed or reported. Collectively, these factors highlight the importance of implementing observational pain assessment tools and routine spinal evaluations in AD patients to detect IVD dysfunction early and prevent the progression to chronic, disabling pathology.

## Conclusion

In this study, we identified significant biomechanical and structural alterations in IVDs of 5xFAD mice, a model of AD, despite structural and compositional measures appearing normal. Specifically, 5xFAD discs exhibited increased volume and viscoelastic damping compared to wild-type controls. These findings suggest an early biomechanical vulnerability potentially driven by amyloid pathology, which standard histological assessments failed to detect.

The observed biomechanical changes underscore a critical implication: individuals with AD may possess heightened susceptibility to spinal dysfunction due to subtle yet impactful mechanical vulnerabilities. This research positions the 5xFAD mouse model as a valuable system for further exploration of neurodegenerative mechanisms and their influence on spinal biomechanics. Importantly, these findings highlight the need for proactive clinical assessment strategies for spinal health in patients with AD, given the propensity for cognitive impairments to obscure pain reporting and delay treatment.

Future research should focus on longitudinal studies that can delineate the progression of mechanical changes in IVDs over time, providing critical temporal context currently lacking from cross-sectional approaches. Additionally, molecular studies should investigate specific amyloidogenic pathways and matrix remodeling mechanisms implicated in disc degeneration, potentially revealing novel therapeutic targets. The incorporation of detailed behavioral assays alongside quantification of spinal amyloid deposition would greatly enhance our understanding of the functional impacts of these biomechanical changes. Such comprehensive approaches will not only illuminate the early pathological processes linking AD and spinal degeneration but also pave the way for interventions aimed at maintaining spinal health and improving quality of life for individuals affected by neurodegenerative diseases.

## Acknowledgements

This work was supported by the NIH R01AR074441 (SYT), S10OD028573, and P30AR074992. Thank you to the Alafi Neuroimaging Lab for Nanozoomer Access (NIH S10RR027552). KAW was supported by T32AG000266 and Buck Catalyst^X^ award from Bob and Alex Griswold. We also acknowledge the support of NIH R01AG061879 (LME) and P01AG066591 (LME).

